## Supplementary material for "Dissecting regulatory pathways for transcription recovery following DNA damage reveals a non-canonical function of the histone chaperone HIRA": Supplemntary figures and legends

### Supplementary Figure 1

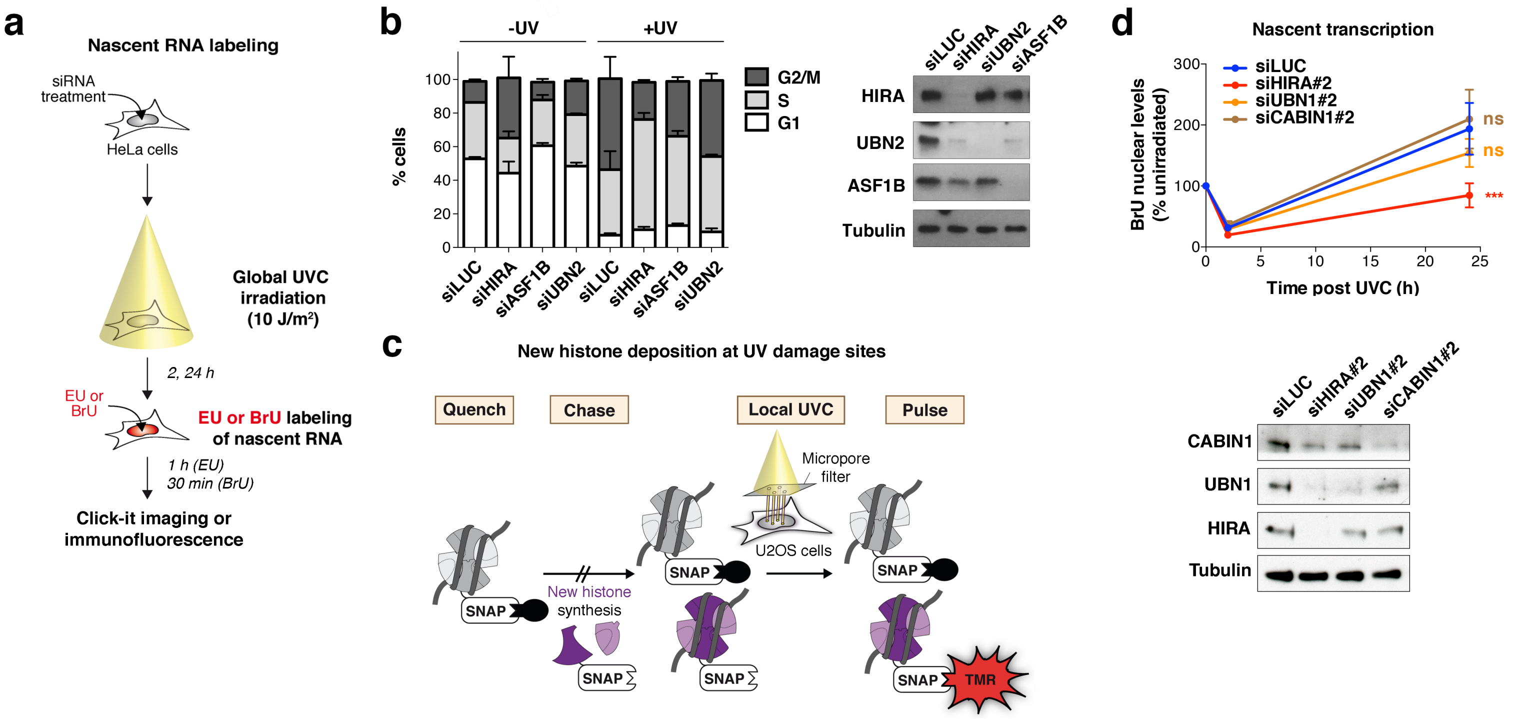

### Supplementary Figure 2

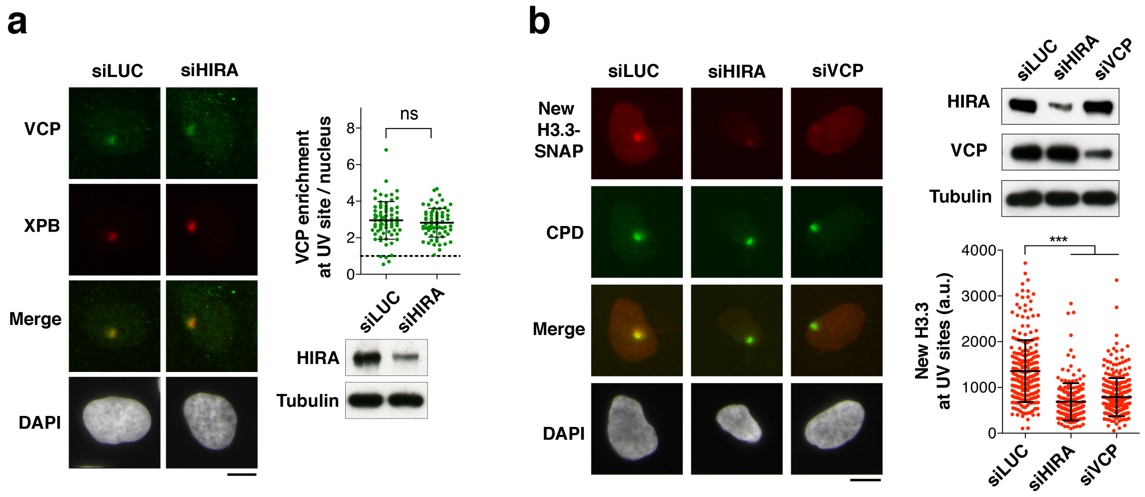

### Supplementary Figure 3

**a**

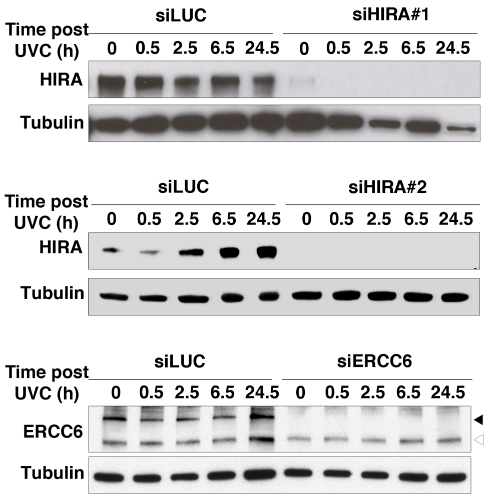


**b**

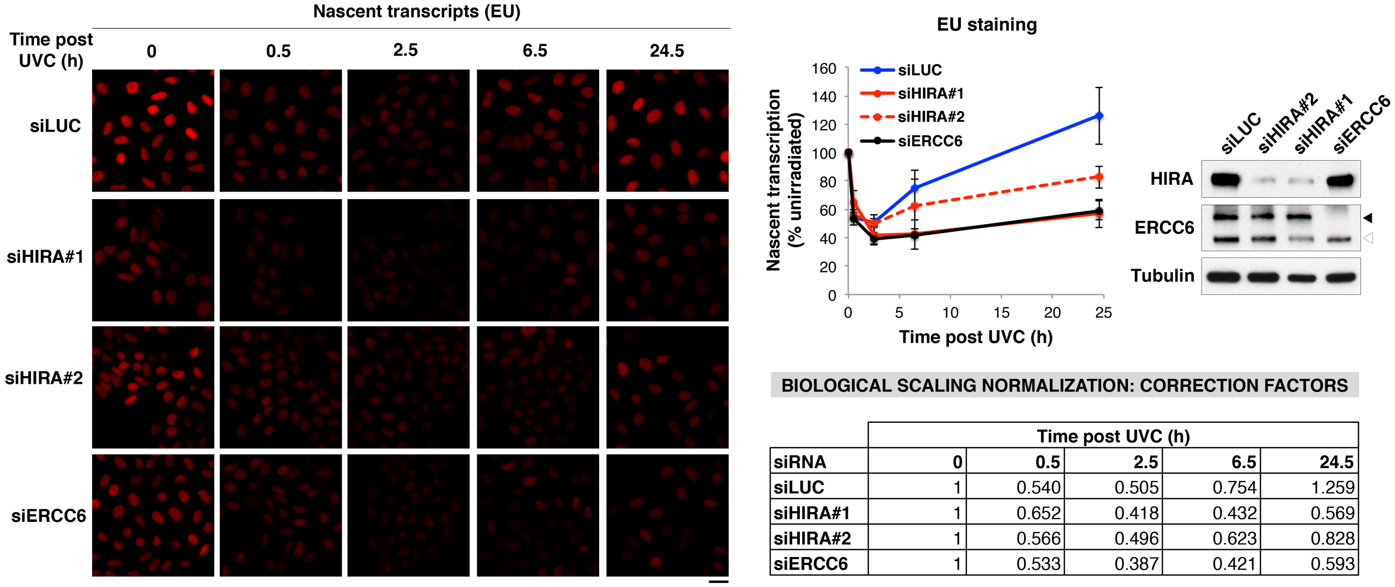


BIOLOGICAL SCALING NORMALIZATION: CORRECTION FACTORS

| | Time post UVC (h) | | | | |
|---|---|---|---|---|---|
| siRNA | 0 | 0.5 | 2.5 | 6.5 | 24.5 |
| siLUC | 1 | 0.540 | 0.505 | 0.754 | 1.259 |
| siHIRA#1 | 1 | 0.652 | 0.418 | 0.432 | 0.569 |
| siHIRA#2 | 1 | 0.566 | 0.496 | 0.623 | 0.828 |
| siERCC6 | 1 | 0.533 | 0.387 | 0.421 | 0.593 |

**c**

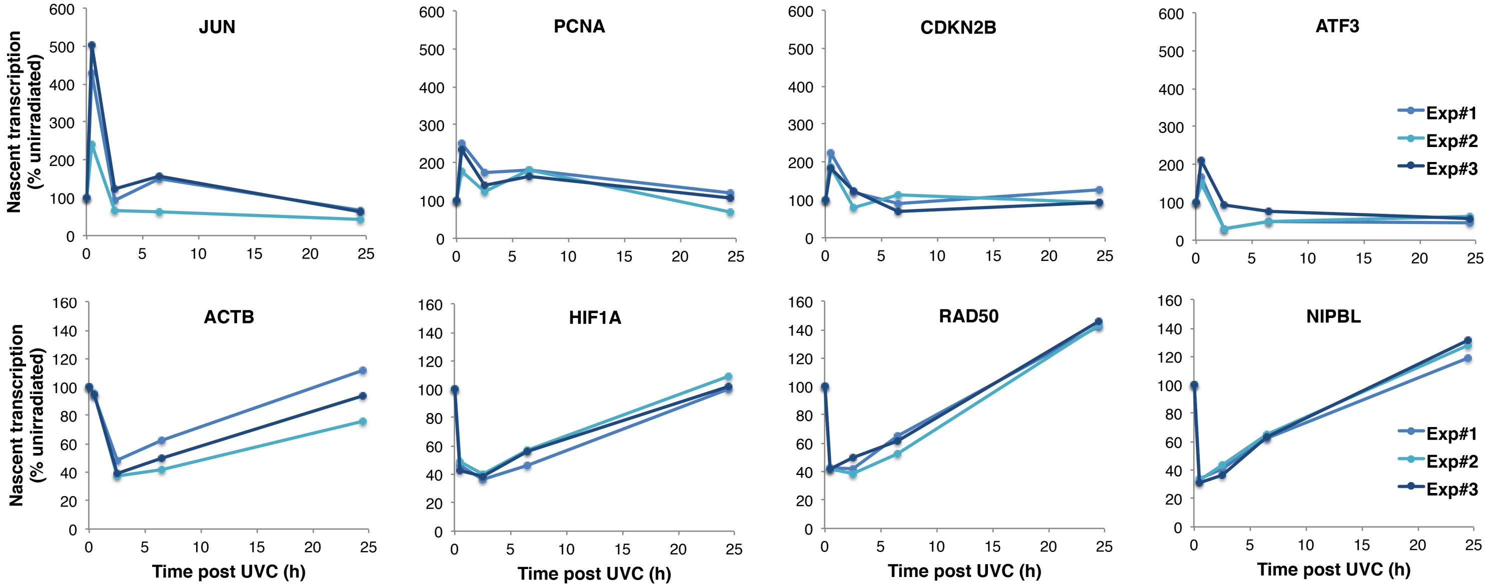


UV-INDUCED GENES

| n | (% expressed genes) | median length |
|---|---|---|
| 702 | 4% | 6.2 kb |
| 425 | 3% | 7.8 kb |
| 759 | 5% | 5.7 kb |

UV-REPRESSED GENES

| n | (% expressed genes) | median length |
|---|---|---|
| 6285 | 38% | 48.7 kb |
| 7395 | 46% | 35.7 kb |
| 6898 | 41% | 40.4 kb |

### Supplementary Figure 4

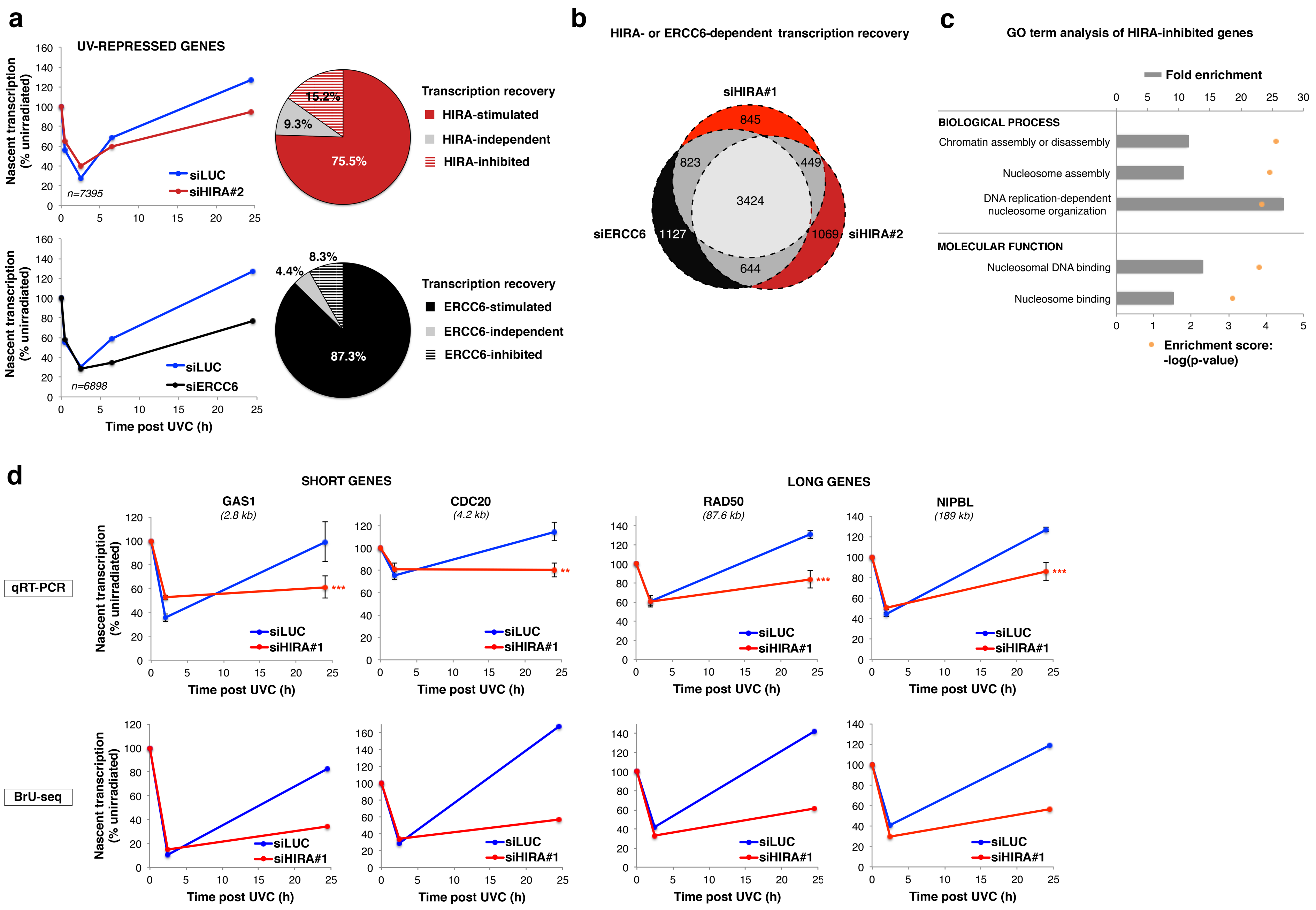

### Supplementary Figure 5

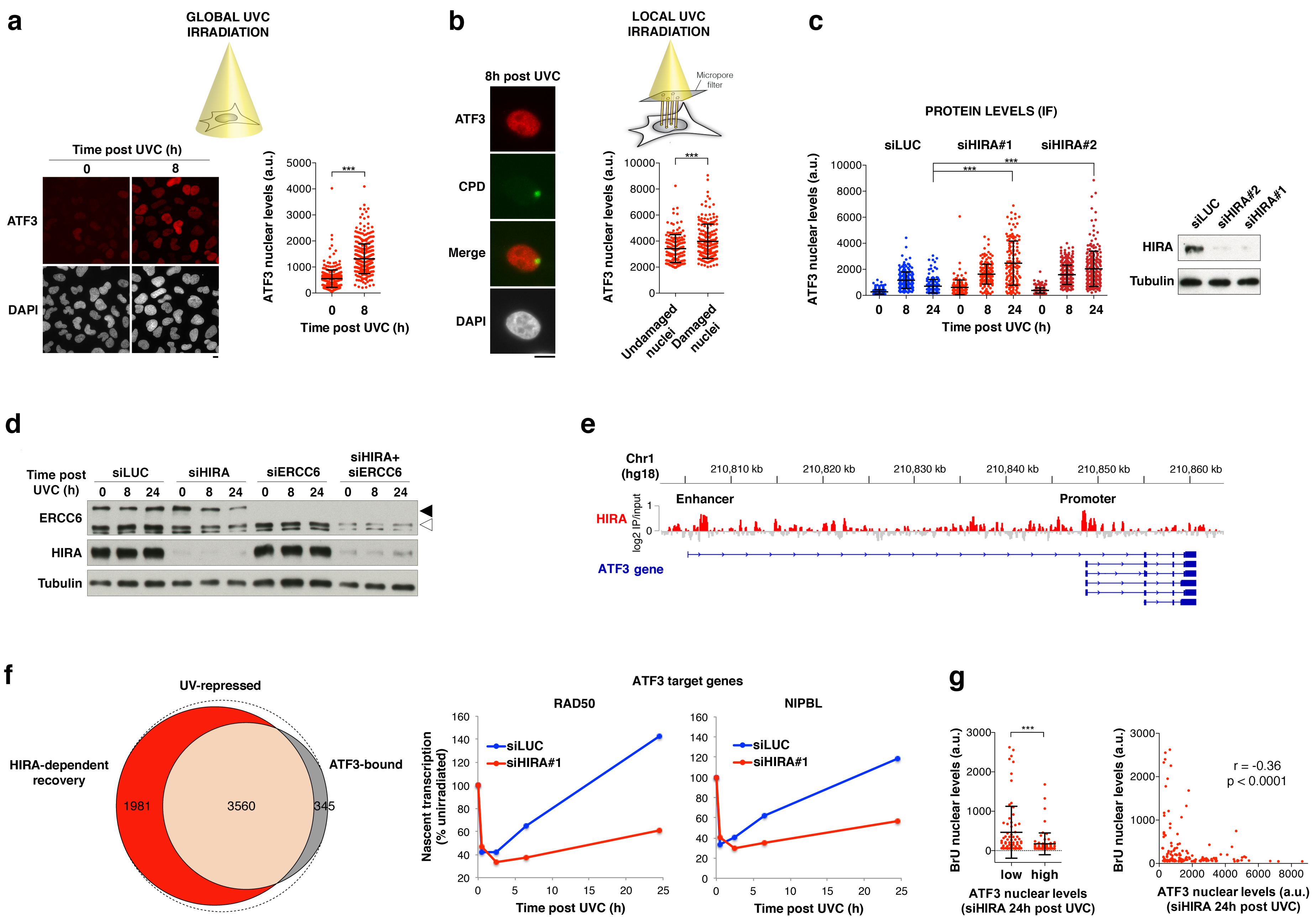

**Supplementary figure legends**

**Supplementary Fig. 1: Functional consequences of HIRA complex subunit knockdown**

**a**, Scheme of the procedure for nascent RNA labeling with EU or BrU in HeLa cells treated with siRNAs. Cells were irradiated at different time points after siRNA transfection to be labeled all at the same time with EU or BrU and harvested right after. **b**, Cell cycle distribution of HeLa cells treated with the indicated siRNAs and analyzed by flow cytometry before (-UV) and 24 h after global UVC irradiation at 10 J/m$^2$ (+UV). Error bars, s.d. from three independent experiments. **c**, Scheme of the assay for monitoring new H3.3 deposition at sites of local UVC irradiation in U2OS H3.3-SNAP cells. Pre-existing SNAP-tagged histones are quenched with a non-fluorescent substrate (block) and histones neosynthesized during the chase period are labeled with the red fluorescent substrate tetramethylrhodamine (TMR)-star during the pulse step. Local UVC irradiation through micropore filters is performed immediately before the pulse step. **d**, Nascent transcript levels analyzed by bromo-uridine (BrU) incorporation in UVC-irradiated HeLa cells treated with the indicated siRNAs. The graphs represent nascent transcript levels relative to unirradiated cells (error bars, s.e.m. from six independent experiments scoring at least 94 cells per condition). siRNA efficiencies are controlled by western-blot (siLUC, control; Tubulin, loading control).

**Supplementary Fig. 2: HIRA and VCP interplay at UVC damage sites**

**a**, VCP recruitment to UVC damage sites (marked by the repair factor XPB) analyzed by immunostaining 5 min after local UVC irradiation in U2OS cells treated with the indicated siRNAs (siLUC, control). **b**, New H3.3 accumulation at UVC damage sites (marked by CPD immunostaining) analyzed 2 h post local UVC irradiation in U2OS H3.3-SNAP cells treated with the indicated siRNAs (siLUC, control). siRNA efficiencies are controlled by western blot (Tubulin, loading control). The scatter plots show data from a representative experiment (bars, mean; error bars, s.d. from at least 67 cells). Similar results were obtained in three independent experiments. Scale bar, 10 µm.

**Supplementary Fig. 3: Biological scaling normalization for Bru-seq and examples of UV-regulated genes**

**a**, Western-blot controls of HIRA and ERCC6 knock-downs at the indicated time points post UVC irradiation in HeLa cells (siLUC, control; black arrowhead, full-length ERCC6; white arrowhead, ERCC6 splice variant; Tubulin, loading control). Each western-blot corresponds to a Bru-seq experiment. **b**, Ethynyl-uridine (EU) labeling of nascent transcripts in HeLa cells treated with the indicated siRNAs (siLUC, control). Scale bar, 10 µm. The western-blot shows the efficiency of protein knock-down by siRNA at the end of the experiment as in (a). The graph represents nascent transcript levels post UVC irradiation normalized to before damage with mean values from 3 independent experiments (error bars, s.d.). These values are used as correction factors for the normalization of Bru-seq data (table). **c**, Examples of UV-induced (top) and UV-repressed genes (bottom) analyzed by Bru-seq with biological scaling normalization (3 independent experiments). Nascent transcript levels post UVC irradiation normalized to before damage are presented on the graphs. Statistics for each category of genes are shown in the tables (n, number of genes).

**Supplementary Fig. 4: Bru-seq analysis and validation by qRT-PCR**

**a**, Bru-seq analysis of UV-repressed genes in HeLa cells treated with the indicated siRNAs (siLUC, control). Nascent transcript levels post UVC irradiation normalized to before damage averaged over n genes are presented on the graphs. The pie charts show UV-repressed genes as a function of their dependency on HIRA or ERCC6 for transcription recovery. **b**, Venn diagrams showing the number of genes requiring HIRA or ERCC6 for transcription recovery 24h30 post UVC irradiation. **c**, Gene ontology analysis of escapees that recover transcription better in the absence of HIRA (HIRA-inhibited genes identified in both experiments with siHIRA#1 and siHIRA#2). **d**, Validation of Bru-seq results by qRT-PCR on short and long genes in HeLa cells treated with the indicated siRNAs (siLUC, control). Bru-seq and qRT-PCR results are presented one above the other for comparison. Error bars represent s.e.m. from three independent experiments.

**Supplementary Fig. 5: HIRA impact on ATF3 levels following UVC irradiation**

**a**, Upregulation of ATF3 analyzed by immunofluorescence in U2OS cells 8 h after global UVC irradiation compared to unirradiated cells. **b**, ATF3 nuclear levels analyzed by immunofluorescence 8 h after local UVC irradiation through micropore filters in U2OS cells. UVC damage sites are revealed by CPD immunodetection. **c**, ATF3 protein levels analyzed by immunofluorescence at the indicated time points post UVC irradiation in HeLa cells treated with the indicated siRNAs (siLUC, control). Knock-down efficiencies are controlled by western-blot (Tubulin, loading control). The scatter plot shows ATF3 levels quantified in cell nuclei (bars, mean; error bars, s.d. from at least 125 cells). Similar results were obtained in two independent experiments. a.u., arbitrary units. **d**, Knock-down efficiencies of data presented in Fig. 6b controlled by western-blot (black arrowhead, ERCC6 full-length; white arrowhead, ERCC6 splice variant; Tubulin, loading control). **e**, HIRA ChIP-seq profile on the ATF3 gene in HeLa cells (data from Pchelintsev et al., 2013; log2 fold change relative to input obtained by bigwigCompare and plotted with Integrative Genomics Viewer). **f**, Venn diagram showing the overlap between UV-repressed genes that require HIRA for transcription recovery and those bound by ATF3 (based on Bru-seq experiment with siHIRA#1 and on ATF3 ChIP-seq data 8 h post UVC from Epanchintsev et al., 2017). The positions of ATF3 ChIP-seq peaks (with FDR<1, fold enrichment >10) were intersected with the positions of the nascent transcripts extended by 5 kb upstream to include promoter regions. The graphs show examples of ATF3-target genes that require HIRA for transcription recovery post UVC (Bru-seq data with biological scaling normalization). **g**, Anti-correlation between ATF3 and BrU nuclear levels in HIRA-knocked down cells 24h post UVC irradiation. Left graph, ATF3 levels were partitioned in two groups based on the median value (bars, mean; error bars, s.d. from 73 cells). Right graph, Spearman correlation. r, correlation coefficient (n=146). Similar results were obtained in three independent experiments. Scale bars, 10 μm.
